# Impaired *in vitro* Interferon-γ production in patients with visceral leishmaniasis is improved by inhibition of PD1/PDL-1 ligation

**DOI:** 10.1101/2021.11.01.466750

**Authors:** Yegnasew Takele, Emebet Adem, Susanne Ursula Franssen, Rebecca Womersley, Myrsini Kaforou, Michael Levin, Ingrid Müller, James Anthony Cotton, Pascale Kropf

## Abstract

Visceral leishmaniasis (VL) is a neglected tropical disease that causes substantial morbidity and mortality and is a growing health problem in Ethiopia, where this study took place. Most individuals infected with *Leishmania donovani* parasites will stay asymptomatic, but some develop VL that, if left untreated, is almost always fatal. This stage of the disease is associated with a profound immunosuppression, characterised by impaired production of Interferonγ (IFNγ), a cytokine that plays a key role in the control of *Leishmania* parasites, and high expression levels of an inhibitory receptor, programmed cell death 1 (PD1) on CD4^+^ T cells. Here, we tested the contribution of the interaction between the immune checkpoint PD1 and its ligand PDL-1 on the impaired production of IFNγ in VL patients. Our results show that in the blood of VL patients, not only CD4^+^, but also CD8^+^ T cells express high levels of PD1 at the time of VL diagnosis. Next, we identified PDL-1 expression on different monocyte subsets and neutrophils and show that PDL-1 levels were significantly increased in VL patients. PD1/PDL-1 inhibition resulted in significantly increased production of IFNγ, suggesting that therapy using immune checkpoint inhibitors might improve disease control in these patients.

## INTRODUCTION

Visceral leishmaniasis (VL) is one of the most neglected tropical diseases [1]. An estimated 550 million individuals are at risk of VL in high-burden countries: 17,082 new cases of VL were reported in 2018, with Brazil, Ethiopia, India, South Sudan and Sudan – which each reported more than 1000 VL cases – represented 83% of all cases globally in that year [2]. The remote location of VL endemic areas and the lack of surveillance make its likely that this is a significant underestimate of the real burden of VL in endemic areas. VL imposes a huge pressure on low and middle income countries and delays economic growth, with an approximate annual loss of 2.3 million disability-adjusted life years [3]. In Ethiopia, VL is caused by *Leishmania* (*L.*) *donovani* and is one of the most significant vector-borne diseases, with over 3.2 million people at risk of infection [4]. VL is a growing health problem, with spreading endemic areas and a steady increase in incidence since 2009 [5].

The majority of infected individuals control the parasite replication and do not progress to disease, they remain asymptomatic. In contrast, some individuals will progress and develop visceral leishmaniasis that is characterised by hepatosplenomegaly, fever, pancytopenia and wasting; this stage of the disease is generally fatal if left untreated [6, 7]. One of the main immunological characteristic of VL patients is their profound immunosuppression [8]: these patients do not respond to the leishmanin skin test, their peripheral blood mononuclear cells (PBMCs) have an impaired capacity to produce IFNγ and proliferate in response to *Leishmania* antigen; this dysfunctional response to antigen challenge is restored following successful chemotherapy [9–11]. These findings show that T cell responses are impaired in VL patients, but the mechanisms leading to this impairment remain to be fully understood Using a whole blood assay (WBA), we have previously shown that whole blood [12] cells from VL patients from Northern Ethiopia displayed an impaired capacity to produce IFNγ in response to stimulation with soluble *Leishmania* antigens (SLA) at time of diagnosis; but that these cells gradually regained their capacity to produce IFNγ over time after successful treatment [13, 14].

We have recently shown that high levels of PD1 on CD4^+^ T cells – an inhibitory receptor that can be expressed on exhausted T cells – was a hallmark of VL patients at time of diagnosis and that this was associated with low production of IFNγ [14]. The interaction of PD1 with its ligand PDL-1 contribute to T cell dysfunction [15]. Therefore, in this study we aimed to identify which cells express PDL-1; and determine whether the impaired production of IFNγ can be improved by interfering with the PD1/PDL1 pathway.

## MATERIAL AND METHODS

### Subjects and sample collection

This study was approved by the Institutional Review Board of the University of Gondar (IRB, reference O/V/P/RCS/05/1572/2017), the National Research Ethics Review Committee (NRERC, reference 310/130/2018) and Imperial College Research Ethics Committee (ICREC 17SM480). Informed written consent was obtained from each patient and control. For this cross-sectional study, 10 healthy male non-endemic controls were recruited among the staff of the University of Gondar, Ethiopia and 16 male patients with visceral leishmaniasis (VL patients) were recruited from the Leishmaniasis Treatment and Research Center, University of Gondar. The exclusion criterion was age <18 years. The diagnosis of VL was based on positive serology (rK39) and the presence of *Leishmania* amastigotes in spleen aspirates [16]. All patients were treated with a combination of sodium stibogluconate (20mg/kg body weight/day), and paromomycin (15mg/kg body weight/day) injections for 17 days according to the Guideline for Diagnosis and Prevention of Leishmaniasis in Ethiopia [17].

### Sample collection and processing

8ml of blood was collected in heparinised tubes and was used as follows: 3ml for the whole blood assay (WBA) and 5ml to purify PBMC and neutrophils as described in [18] for flow cytometry.

The WBA was performed as described in [13]. For the blockade of PD1/PDL-1 interactions, whole blood [12] cells were stimulated with SLA in the presence or in the absence of 1μg of anti-PD-1 (Nivolumab) Humanized Antibody (BioVision). IFNγ levels were measured in the supernatant of the WBA using IFN gamma ELISA Kit (Invitrogen) according to the manufacturer’s instructions.

Flow cytometry: the following antibodies were used: CD4^FITC^, CD8^PE^ ^CY7^, PD1^PE^, PDL-1^PE^, CD15^APC^ (eBioscience), CD14^APC^ and CD16^FITC^ (Biolegend) as described in [14].

Acquisition was performed using a BD Accuri™ C6 flow cytometer and data were analysed using BD Accuri C6 analysis software.

### Statistical analysis

Data were evaluated for statistical differences as specified in the legend of each figure. The following tests were used: Mann-Whitney or Wilcoxon tests. Differences were considered statistically significant at *p*<0.05. Unless otherwise specified, results are expressed as median±SEM. *=*p*<0.05, **=*p*<0.01, ***=*p*<0.001 and ****=*p*<0.0001.

## RESULTS

### Clinical and haematological parameters

The cohort of 16 VL patients and 10 healthy non-endemic controls that were recruited for this study were all male and aged-matched (Table 1). *Leishmania* amastigotes were present in all splenic aspirates from VL patients (parasite grade (+): 2.5±0.5). All VL patients presented with low BMI, fever, splenomegaly, and pancytopenia (Table 1).

**Table 1:** Clinical and haematological parameters.

|  | **ToD** | **Controls** | **p values** |
| --- | --- | --- | --- |
| **Age (years)** | 25±2.1 | 26±2.2 | 0.5057 |
| **Parasite grade (+)** | 2.5±0.5 | nd | na |
| **BMI (kg/m<sup>2</sup>)** | 16.2±0.4 | 21.0±1.0 | <0.0001 |
| **Fever (°C)** | 37.4±0.2 | 36.0±0.1 | <0.0001 |
| **Spleen size (cm)** | 8.5±1.0 | 0.0±0.0 | <0.0001 |
| **WBC (µl of blood)** | 1.8±0.1×10<sup>3</sup> | 5.8±0.7×10<sup>3</sup> | <0.0001 |
| **RBC (µl of blood)** | 3.0±0.2×10<sup>6</sup> | 5.4±0.2×10<sup>6</sup> | <0.0001 |
| **Platelets (µl of blood)** | 5.7±1.2×10<sup>4</sup> | 24.8±2.0×10<sup>4</sup> | <0.0001 |

A cohort of VL patients (n=16) were recruited at time of diagnosis. Clinical and haematological parameters were assessed and compared to those of a cohort of age- and sex-matched non-endemic healthy controls (n=10). Spleen sizes were measured below the left costal margin. nd= not done, na=not applicable.

### PD1 expression on T cells

Efficient effector functions of specific T cells are of crucial importance for the clinical outcome of visceral leishmaniasis. One of the main immunological features of VL patients in Ethiopia is their impaired ability at time of diagnosis (ToD) to produce antigen-specific IFNγ in a whole blood assay (WBA); after successful treatment the impaired IFNγ production is restored over time [13, 14]. Our recent work [14] showed that the gradual increase in antigen-specific IFNγ production during follow-up is accompanied by a gradual decrease of PD1 expression on CD4^+^ T cells. Results presented in Figures 1A and B show that not only CD4^+^ T cells, but also CD8^+^ T cells expressed PD1. In both T cell subsets, the expression levels of PD1 on CD4^+^ T cells and CD8^+^ T cells were significantly higher as compared to controls (*p*<0.0001 and 0.0004, respectively).

**Figure 1:**
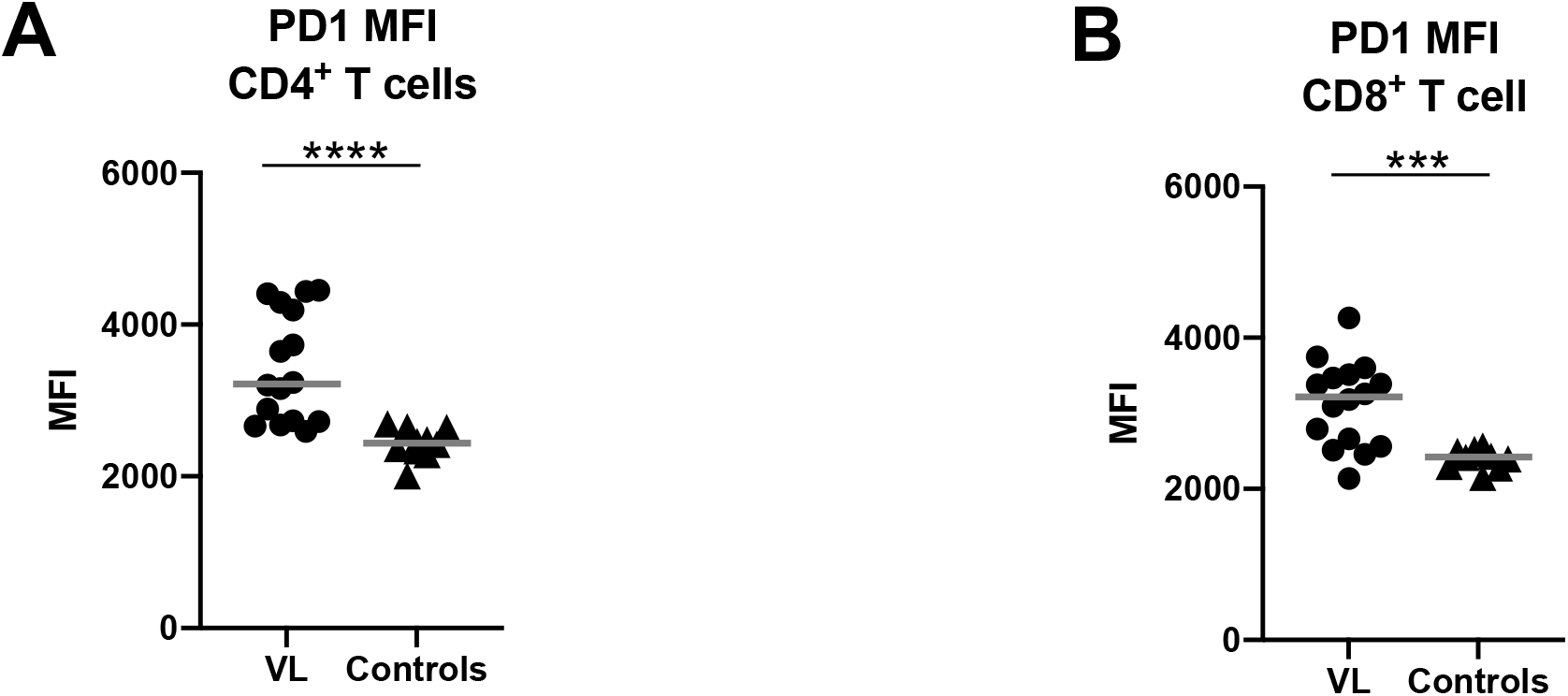
Expression of PD1 on T cells. The Median Fluorescence Intensity [32] of PD1 was measured by flow cytometry on CD4^+^ T cells (**A**) and CD8^+^ T cells (**B**) isolated from VL patients (n=16) and controls (n=10). Statistical differences were determined by a Mann-Whitney test. Each symbol represents the value for one individual, the straight lines represent the median.

### Monocytes and neutrophils express PDL-1

We have hypothesised that the high levels of PD1, via its interaction with PDL-1, is a major contributor of T cell hyporesponsiveness in VL patients [14]. However, the phenotypes of PDL-1 expressing cells have not yet been identified in VL patients. Results presented in Figure 2 show that monocytes in the PBMCs of VL patients express PDL1. All three monocyte subsets: classical (CD14^+^/CD16^low^, Figure 2A), intermediate (CD14^+^/CD16^+^, Figure 2B) and non-classical (CD14^low^/CD16^+^, Figure 2C) expressed significantly more PDL-1 at ToD compared to controls (*p*>0.0001, *p*>0.0001 and *p*=0.0061, respectively; Tables 1A, B and C). The intermediate subsets of monocytes expressed the highest levels of PDL-1 (Figures 2 D and E and Table 2). There was no significant difference in PDL-1 expression between the three subsets of monocytes in the controls (Figure 2E).

**Figure 2:**
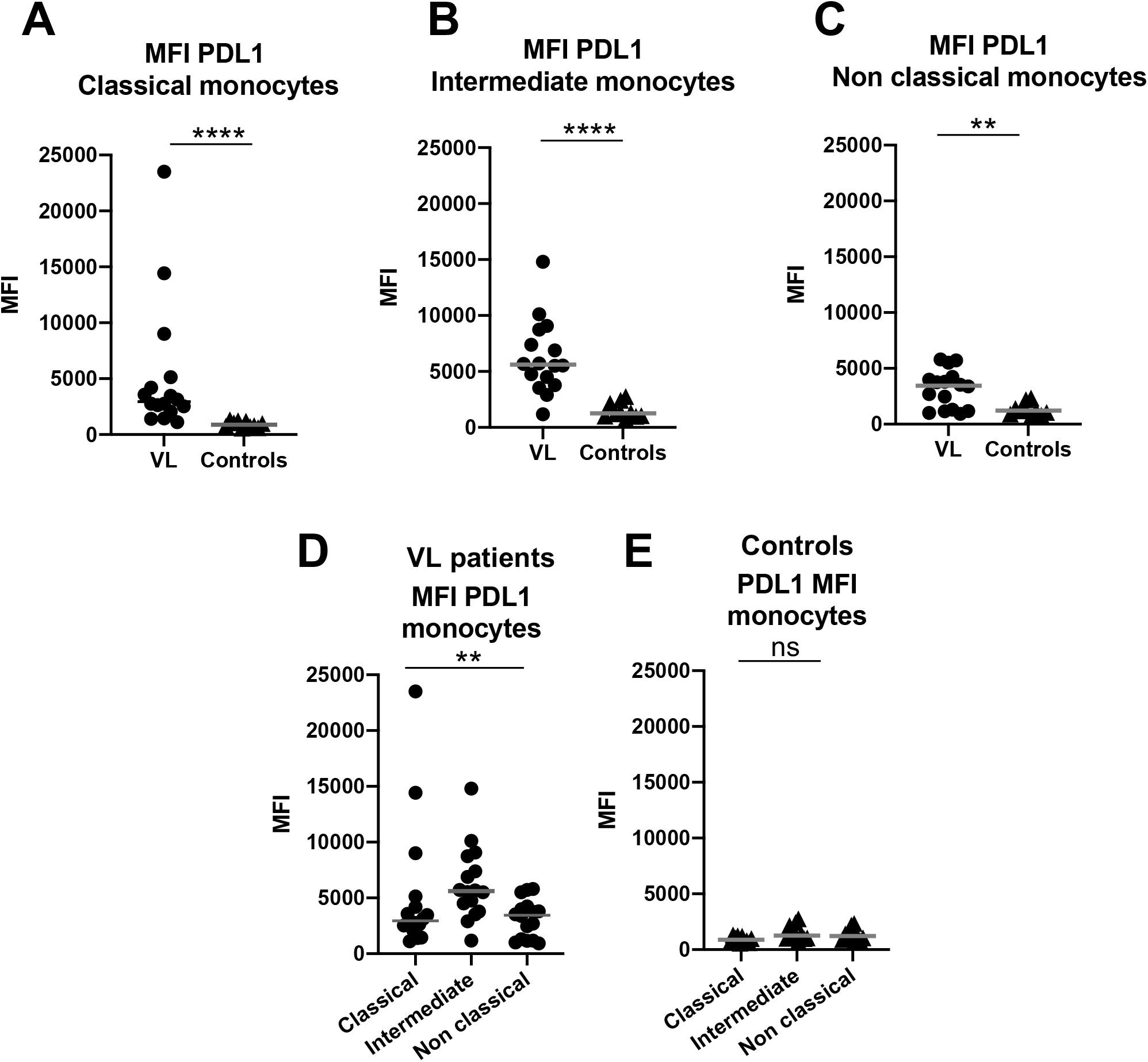
Monocytes express PDL-1. PDL-1 MFI was measured by flow cytometry on monocytes from PBMCs of VL patients n=16) and controls (n=10). **A.** PDL-1 expression on classical monocytes. **B.** PDL-1 expression on intermediate monocytes. **C.** PDL-1 expression on non classical monocytes. Statistical differences were determined by a Mann-Whitney test. Comparison of PDL-1 expression between the 3 different subsets of monocytes from VL patients (**D**) and controls (**E**). Statistical differences between the 3 different subsets were determined by a Kruskal-Wallis test. Each symbol represents the value for one individual, the straight lines represent the median

Next, we assessed if neutrophils also expressed PDL-1 at ToD. Results presented in Figure 3 show that neutrophils expressed significantly higher levels of PDL-1 at ToD than controls (*p*=0.0022).

**Figure 3:**
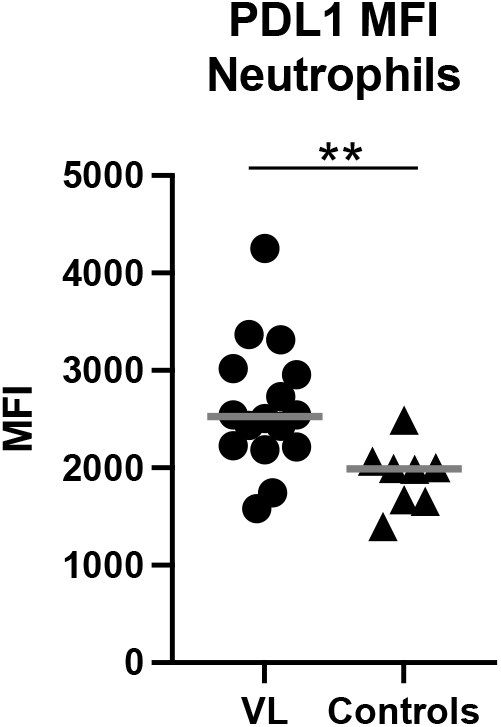
Neutrophils express PDL-1. PDL-1 MFI was measured by flow cytometry on neutrophils in the PBMCs from VL patients (n=16) and controls (n=8). Statistical differences were determined by a Mann-Whitney test. Each symbol represents the value for one individual, the straight lines represent the median

### Interfering with the PD1/PDL-1 pathway results in increased production of IFNγ

Based on the increased expression levels of PD1 on T cells and PDL-1 on monocytes and neutrophils and the low levels of IFNγ produced in the WBA [14], we tested if blockade of the PD1/PDL-1 interaction improves IFNγ production. Results presented in Figure 4 show that interfering with the PD1/PDL-1 ligation resulted in significantly higher levels of IFNγ (p=0.0006), suggesting that the impaired ability of WB cells to produce IFNγ efficiently can be improved by PD1/PDL-1 blockade.

**Figure 4:**
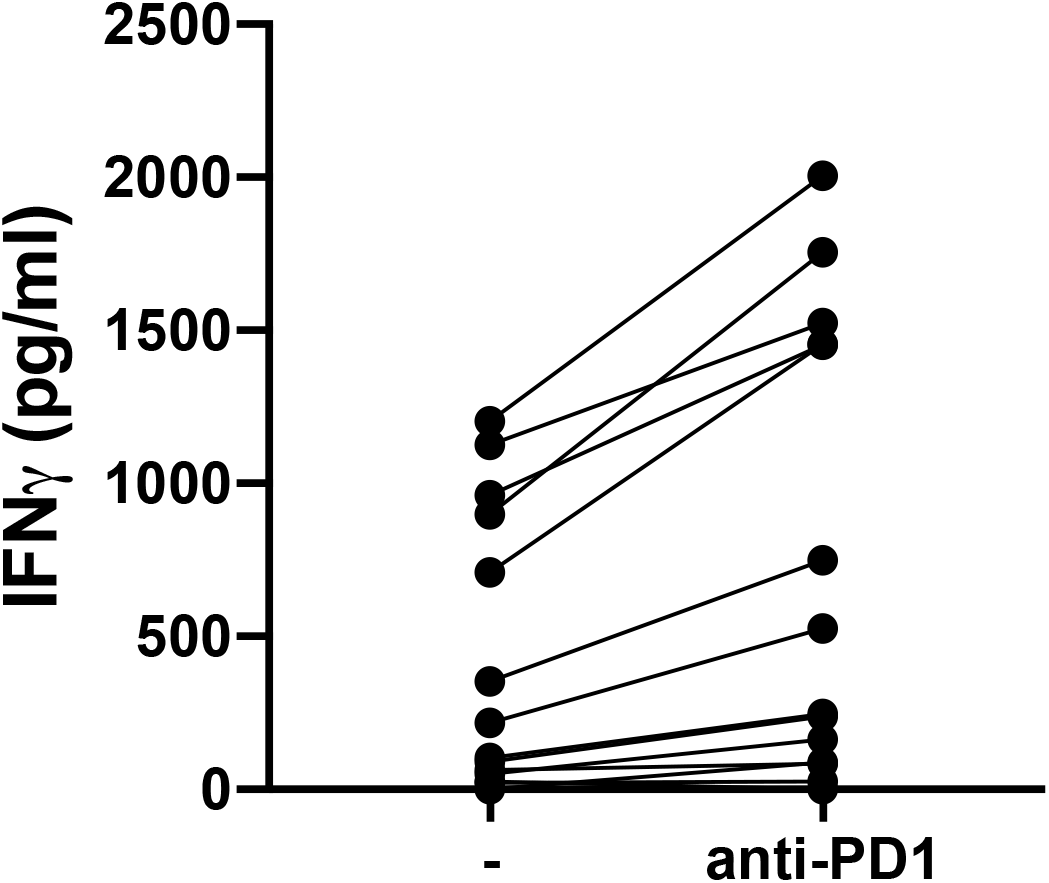
Interfering with the PD1/PDL-1 pathway results in increased production of IFNγ. Whole blood cells from VL patients at ToD (n=8) were stimulated with SLA in the presence (1μg) or absence of anti-PD-1 mAb. IFNγ was measured by ELISA in the supernatant after 24hrs. Statistical differences were determined by a Wilcoxon test. Each symbol represents the value for one individual.

## DISCUSSION

Severe immune suppression has been previously documented in VL patients [9], however, it is still poorly understood. Here we show that at the time of diagnosis of VL both CD4^+^ and CD8^+^ T cells express high levels of PD1. The interaction of inhibitory receptors such as PD1 with their ligand play a crucial role in controlling autoreactivity and immunopathology. These receptors are also upregulated during T cell activation, but this is transient. In contrast, chronic stimulation of T cells results in the maintenance of high levels of PD1 expression on T cells; the duration and degree of this chronic stimulation are key to T cell exhaustion and dysfunction [19]. VL patients present at the Leishmaniasis Treatment and Research Center with severe disease and on average have had VL symptoms for around 2 months [14]. It is therefore likely that the chronic stimulation by *Leishmania* antigen plays a major role in the maintenance of high expression levels of PD1 on T cells in these patients. Several inflammatory mediators have been shown to result in the upregulation of PDL-1 on different cell types such as neutrophils and monocytes [20]. It is well established that high levels of inflammation are common in VL patients [9] and indeed, we found high levels of TNFα, IL-6, IL-8, IFNγ [14], as well as IL-17 [21] in the plasma of VL patients at the time of diagnosis. IL-10 has also been shown to upregulate PDL-1 on monocytes [22] and indeed, high levels of IL-10 in plasma are a hallmark of VL patients [9, 14]. Therefore, we propose that in VL patients, the chronic inflammation and antigenic stimulation results in the upregulation of both PD1 and PDL-1, that results in T cell exhaustion; that is manifested in the whole blood assay by impaired production of IFNγ. Exhausted T cells become hypofunctional, they maintain some of their effector functions such proliferation [15, 19, 23]. Exhausted T cells have been shown to contribute to the control of chronic infections and limit immunopathology [15, 19, 23]. It is therefore possible that during the acute state of infection, exhausted T cells might still contribute to the control of parasite replication, as well as limit tissue damage.

Our results that show that interfering with the PD1/PDL-1 pathway results in increased production of IFNγ disagree with the results presented by Gautam *et al.* [24]. This study was performed in India, where VL patients present at a significantly earlier stage of the disease, with less severe symptoms; and indeed, the results of their study show that the levels of IFNγ measured in the whole blood assay were not impaired at time of diagnosis as compared to those detected after successful treatment [25]. As previously discussed, [13, 14], we propose that the impaired production of IFNγ we measured in the WBA from the Ethiopian patients is closely related to the fact that they present late, often in a critical state.

In Ethiopia, the first line of treatment against visceral leishmaniasis is a combination therapy of Sodium Stibogluconate (SSG) and Paromomycin [17]. While this treatment shows a good safety profile and a good efficacy, there are still severe side effects, such as cardiotoxicity and nephrotoxicity [26]. In our latest study, four individuals from our cohort of 50 VL patients died during treatment [14]. PD1/PDL-1 blockade has emerged as a front-line treatment for several types of cancer [27].

However, little is known about its potential use in chronic infectious diseases. In a non-human primate model of simian immunodeficiency virus blockade of the PD-1 pathway improved T cell effector functions and resulted in more efficient viral control [28]. Paradoxically, in a 3D cell culture model of tuberculosis despite the increased levels of both PD1 and PDL-1, blockade of PD1 promoted the replication of *M. tuberculosis* [29]. Several studies in experimental models have suggested that immune checkpoint blockade maybe relevant to treat several infectious diseases [30]. Even though immune checkpoint blockade can cause organ-specific immune-related adverse events, such as hepatitis and colitis, as well as systemic inflammation [31]; further studies are needed to determine whether blockade of the PD-1/PDL1 pathway can be used to improve therapies against infectious diseases. In the case of visceral leishmaniasis, it might be particularly useful in tackling the severe form of the disease, to prevent the treatments’ adverse side effects, by allowing the use of shorter courses or reduced doses of current anti-leishmanial treatments.

## AUTHOR CONTRIBUTIONS

All authors discussed the results and contributed to the final manuscript. Study conception and design: PK, YT, IM; Acquisition of data: YT, EA, PK; Analysis and interpretation of data: YT, EA, SF, RW, MK, ML, IM, JAC, PK; Drafting of manuscript: PK, YT, JAC, IM

## ACKNOWLEDGMENTS

We are grateful to the staff of the Leishmaniasis Research and Treatment Centre for their support and DNDi for supporting the VL treatment service at the University of Gondar. YT is funded by a Wellcome Trust Training Fellowship in Public Health and Tropical Medicine (204797/Z/16/Z). JAC is funded by Wellcome via core funding of the Wellcome Sanger Institute (grant 206194). MK is funded by a Wellcome Trust Sir Henry Wellcome Fellowship (206508/Z/17/Z).

## Notes

<u>Conflict of interest statement</u> The authors have declared that no conflict of interest exists.

### Competing Interest Statement

The authors have declared no competing interest.

